# Group-level progressive alterations in brain connectivity patterns revealed by diffusion-tensor brain networks across severity stages in Alzheimer’s disease

**DOI:** 10.1101/105270

**Authors:** J. Rasero, C. Alonso-Montes, I. Diez, L. Olabarrieta-Landa, L. Remaki, I. Escudero, B. Mateos, P. Bonifazi, M. Fernandez, J.C. Arango-Lasprilla, S. Stramaglia, J.M. Cortes, for the Alzheimer's Disease Neuroimaging Initiative

**Affiliations:** Dipartamento di Fisica, Universita degli Studi di Bari and INFN. Bari, Italy; Biocruces Health Research Institute. Barakaldo, Spain; BCAM - Basque Center for Applied Mathematics. Bilbao, Spain; Department of Education and Psychology, University of Deusto. Bilbao, Spain; Radiology Service. Cruces University Hospital. Barakaldo, Spain; IKERBASQUE: The Basque Foundation for Science. Bilbao, Spain; Neurology Service. Cruces University Hospital. Barakaldo, Spain; Department of Cell Biology and Histology. University of the Basque Country. Leioa, Spain

**Keywords:** Diffusion-Tensor Imaging, Brain networks, Alzheimer’s disease, Severity progression

## Abstract

Alzheimer’s disease (AD) is a chronically progressive neurodegenerative disease highly correlated to aging. Whether AD originates by targeting a localized brain area and propagates to the rest of the brain across disease-severity progression is a question with an unknown answer. Here, we aim to provide an answer to this question at the group-level by looking at differences in diffusion-tensor brain networks. In particular, making use of data from Alzheimer's Disease Neuroimaging Initiative (ADNI), four different groups were defined (all of them matched by age, sex and education level): *G*_1_ (*N*_1_=36, healthy control subjects, Control), *G*_2_ (*N*_2_=36, early mild cognitive impairment, EMCI), *G*_3_ (*N*_3_=36, late mild cognitive impairment, LMCI) and *G*_4_ (*N*_4_=36, AD). Diffusion-tensor brain networks were compared across three disease stages: stage I 3(Control vs EMCI), stage II (Control vs LMCI) and stage III (Control vs AD). The group comparison was performed using the multivariate distance matrix regression analysis, a technique that was born in genomics and was recently proposed to handle brain functional networks, but here applied to diffusion-tensor data. The results were three-fold: First, no significant differences were found in stage I. Second, significant differences were found in stage II in the connectivity pattern of a subnetwork strongly associated to memory function (including part of the hippocampus, amygdala, entorhinal cortex, fusiform gyrus, inferior and middle temporal gyrus, parahippocampal gyrus and temporal pole). Third, a widespread disconnection across the entire AD brain was found in stage III, affecting more strongly the same memory subnetwork appearing in stage II, plus the other new subnetworks,including the default mode network, medial visual network, frontoparietal regions and striatum. Our results are consistent with a scenario where progressive alterations of connectivity arise as the disease severity increases and provide the brain areas possibly involved in such a degenerative process. Further studies applying the same strategy to longitudinal data are needed to fully confirm this scenario.

## 1. Introduction

Alzheimer's disease (AD), the most common form of dementia, is a chronically progressive neurodegenerative disease highly correlated to aging; indeed, although the prevalence of clinically manifested AD is about 2% at the age of 65 years, it increases to 30% at the age of 85 years (Wimo et al. 1997).

AD is characterized by an accumulation of beta-amyloid plaques and neurofibrillary tangles composed of tau amyloid fibrils (Hardy 2006) associated with synapse loss and neurodegeneration leading to long-term memory impairment and other cognitive problems. To date, there is no treatment known to slow down the progression of this disorder.

The initial AD pathology develops many years before the cognitive and functional impairments are evident. Different terms have been used to describe this disease-starting condition, including pre-dementia and prodromal AD and, more often, MCI (mild cognitive impairment). The concept of MCI varied over the past two decades and has been classified into different broad categories depending on memory performance and the number of impaired cognitive functions (Mueller *et al.* 2005).

An accurate prediction for the conversion from MCI to AD can help to clinicians to evaluate AD risk pre-symptomatically, initiate treatments at early stage, and monitor their effectiveness (Cheng *et al.* 2015, Li *et al.* 2014). However, such a prediction is challenging, as the MCI group is highly heterogeneous and only a few patients convert to AD, a rate of about 8% to 15% convert per year (Ritter *et al.* 2015, Mitchell and Shiri-Feshki 2009). However, the amnestic subtype of MCI is more prevalent than the non-amnestic MCI (Petersen *et al.* 2010), and has an annual conversion rate higher of about 30% to 40% (Schmidtke and Hermeneit 2008, Rozzini *et al.* 2007, Geslani *et al.* 2005).

This study aims to search for neuroimaging biomarkers that can account for differences with respect to a healthy control population from the early to the final stages of AD. Multitude of different neuroimaging studies has addressed the conversion from MCI to AD, see (Zhang *et al.* 2014) and references therein. In relation to structural magnetic resonance imaging (MRI), it was shown that the hippocampus volume and the volume from other subcortical structures at MCI were well correlated to a worse progression to AD, with accuracy of about 65% in the prediction from MCI to AD (Teipel *et al.* 2015).

Rather than assuming that specific brain regions are affected in AD, a blind approach using multiple regions of interest has been shown to achieve a better predictive accuracy (of about 80%) of the conversion from MCI to AD (Westman *et al.* 2011, Eskildsen *et al.* 2013, Liu *et al.* 2013). The use of tensor diffusion MRI in combination with structural MRI has provided better results as compared to only structural MRI, showing that white-matter integrity of the fornix, cingulum, and parahippocampal gyrus provided accuracy varying from 80% to even 95% (Wee *et al.* 2013, Mielke *et al.* 2012, Douaud *et al.* 2013).

Initiatives like the Alzheimer's Disease Neuroimaging Initiative (ADNI) provide important resources to study AD to the research community (including demographic data, imaging datasets, cognitive 1 tests, etc.), pushing forward multimodal studies correlating different imaging modalities to neuropsychological functioning. Interestingly, ADNI also allows the possibility of studying variations in the images at a group level across disease's progression, as brain images are categorized in different groups ranging from Control to AD, with two intermediate stages, early and late mild cognitive impairment, EMCI and LMCI, respectively. Importantly, although EMCI and LMCI patients have memory impairment (Medina *et al.* 2006), the conversion rate to AD is only between 8-15% per year (Mitchell and Shiri-Feshki 2009), making this group have a special relevance in the development of novel imaging techniques that could correlate with disease progression.

Despite extensive research shedding light into the MCI to AD conversion, the precise mechanisms and clinical variables responsible for such progression are poorly understood, mainly due to the lack of time-resolved longitudinal studies in large populations. Taking into consideration previous work (Khedher *et al.* 2015, Douaud *et al.* 2011, Bosch *et al.* 2012, Liu *et al.* 2013, Acosta-Cabronero *et al.* 2012, Preti *et al.* 2012), the present study focus on the variations of brain networks across AD progression at a group level. It is hypothesized that if in the transition from Control to MCI the connectivity pattern of some subnetworks are altered, in further disease stages the alterations of the same subnetworks will coexist together with alterations of new different subnetworks in the AD brain, in a manner that connectivity alterations will finally extend to the rest of the brain.

## 2. Material and Methods

### 2.1 Ethics

The present study made use of ADNI data previously collected in 50 different institutions. Participants provided informed consent before recruitment and data collection started. In addition, participants filled questionnaires approved by each participating site’s Institutional Review Board (IRB). The complete list of ADNI sites’ IRBs can be found using the following link: http://adni.loni.ucla.edu/about/data-statistics/.

### 2.2 Alzheimer's Disease Neuroimaging Initiative (ADNI)

Diffusion tensor imaging (DTI) data was used in this paper from ADNI database http://adni.loni.usc.edu. ADNI was launched in 2003 by the Nat. Inst. on Aging (NIA), the Nat. Inst. Biomedical Imaging and Bioengineering (NIBIB), the Food and Drug Administration (FDA), private pharmaceutical companies and non-profit organizations, as a $60 million, 5-year public-private partnership. ADNI's main goal has been to test whether serial MRI, positron emission tomography (PET), other biological markers, and clinical and neuropsychological assessment can be combined to measure the progression of MCI and early AD. Determination of sensitive and specific markers of very early AD progression is intended to aid researchers and clinicians to develop new treatments and monitor their effectiveness, as well as to lessen the time and cost of clinical trials. The Principal Investigator of this initiative is Michael W. Weiner, MD, VA Medical Center and Univ. California – San Francisco. ADNI subjects have been recruited from over 50 sites across the U.S. and Canada. Currently, around 1500 adults were recruited in the different ADNI initiatives, ages 55 to 90,consisting of cognitively normal older (NC), early/late MCI (EMCI/LMCI), significant memory concern (SMC) and early AD (AD) individuals. The follow up duration of each group is specified in the protocols for ADNI-1, ADNI-2 and ADNI-GO, see further information in www.adni-info.org.

### 2.3 Demographic Data

A total number of N=144 subjects were used in this study (Table S1). This number was chosen in order to get the biggest four groups as possible (Control, EMCI, LMCI and AD), balanced by size, age and sex. DTI images were selected and downloaded from ADNI database, belonging to four different groups: Control (N_1_=36), EMCI (N_2_=36), LMCI (N_3_=36) and AD (N_4_=36). Age and sex were balanced across groups (Table 1), respectively, using a t-test and chi-squared test. In addition, it is important to remark that the “years of education” variable was already controlled by the ADNI group classification, for details see Inclusion criteria in page 31 of https://adni.loni.usc.edu/wp-content/uploads/2008/07/adni2-procedures-manual.pdf

**Table 1:**
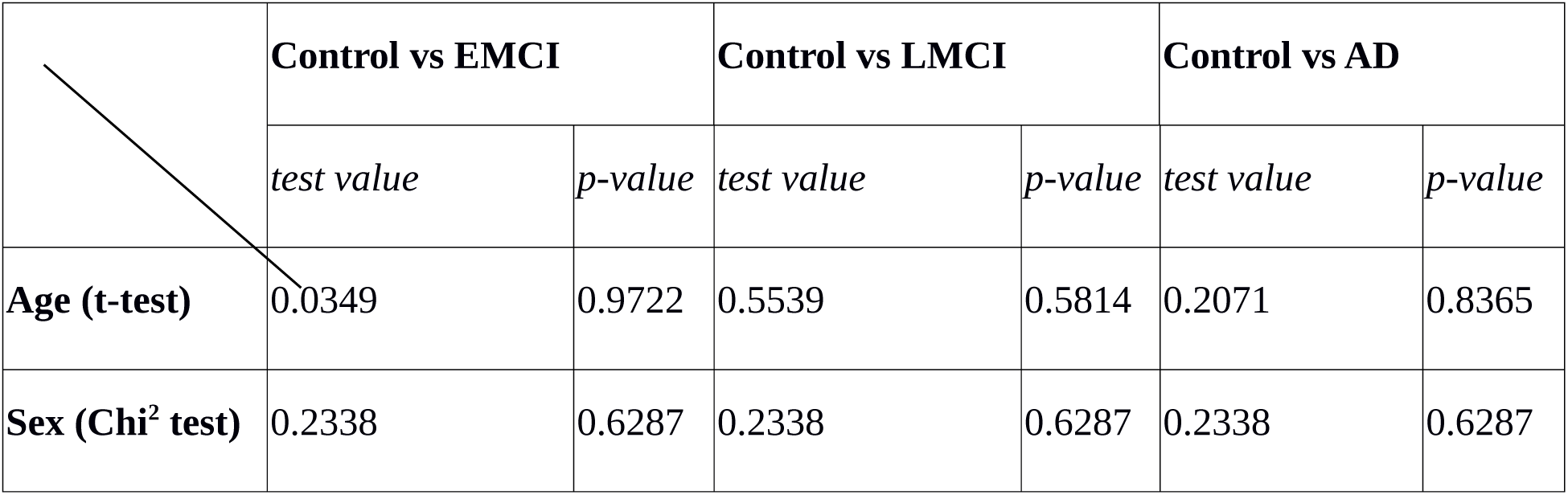
t-test and Chi^2^ test across groups. EMCI: Early mild cognitive impairment; LMCI=Late mild cognitive impairment; AD= Alzheimer disease.

|  | Control vs EMCI |  | Control vs LMCI |  | Control vs AD |  |
| --- | --- | --- | --- | --- | --- | --- |
|  | *test value* | *p-value* | *test value* | *p-value* | *test value* | *p-value* |
| **Age (t-test)** | 0.0349 | 0.9722 | 0.5539 | 0.5814 | 0.2071 | 0.8365 |
| **Sex (Chi<sup>2</sup> test)** | 0.2338 | 0.6287 | 0.2338 | 0.6287 | 0.2338 | 0.6287 |

### 2.4 ADNI group classification

The group labels Control, EMCI, LMCI and AD are based on several test scores, such as the Logical Memory II subscale (LMIIS) from the Wechsler Memory Scale, the Mini-Mental State Examination (MMSE) and the Clinical Dementia Rating (CDR), as well as National Institute of Neurological and Communicative Disorders and Stroke and the <u>AD and Related Disorders Association</u> (NINCDS/ADRDA) criteria in AD cases. In the procedures manual each of the criteria are cited (http://adni.loni.usc.edu/wp-content/uploads/2008/07/adni2-procedures-manual.pdf).

Control subjects are free of memory complaints (beyond normal ageing), verified by a study partner. EMCI, LMCI and AD must have a subjective memory concern as reported by the subject, study partner, or clinician. Details of specific groups are given in Table 2.

**Table 2:**
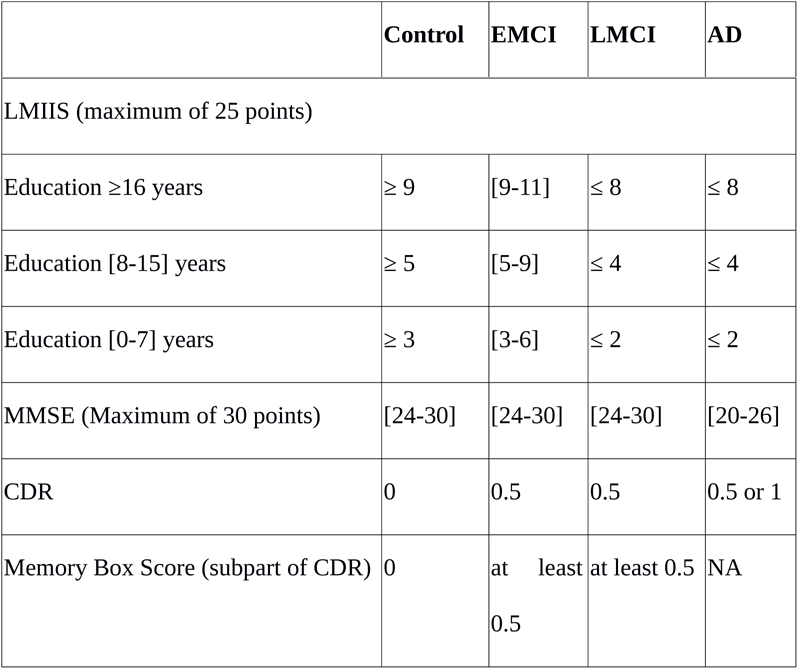
Further information about ADNI group classification. EMCI: Early mild cognitive impairment; LMCI=Late mild cognitive impairment; AD= Alzheimer disease; LMIIS=Logical Memory II subscale; MMSE= Mini Mental State Examination; CDR= Clinical Dementia Rating.

### 2.5 Group-level stages for AD progression

AD progression was defined by three different stages: stage I (control vs EMCI), stage II (control vs LMCI) and stage III (control vs AD). Further details are given in Figure 1.

**Figure 1:**
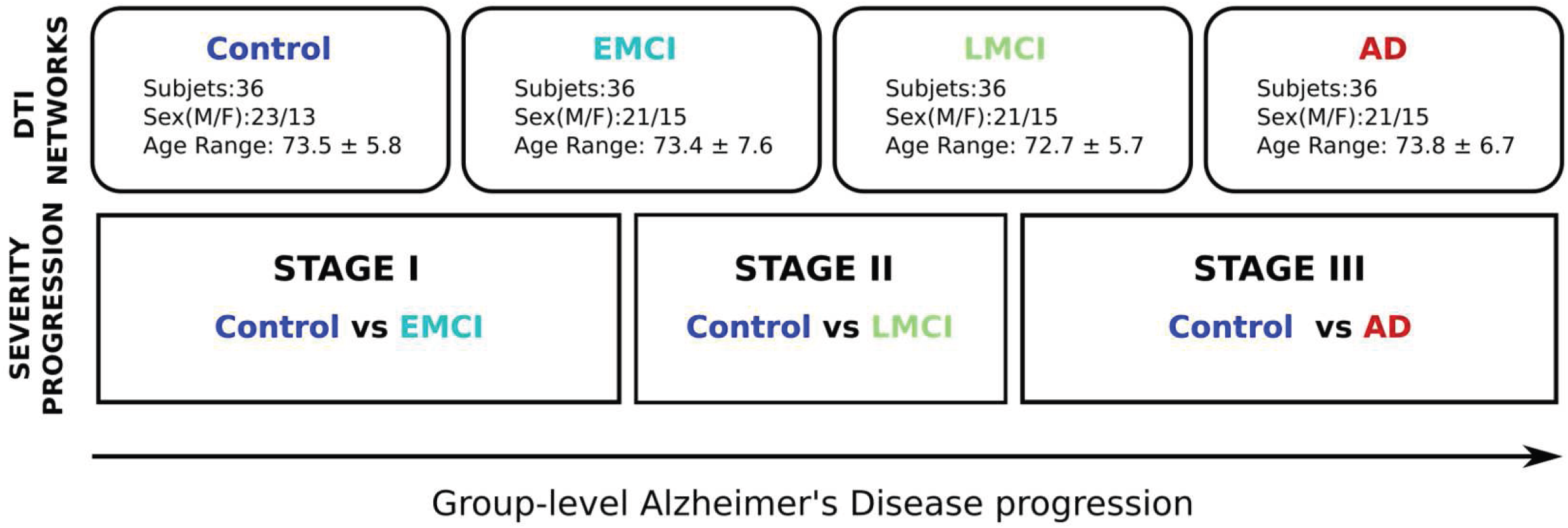
Methodological sketch. Alzheimer’s disease progression is addressed across three stages. Four groups of 36 subjects each at different stages of AD (Control, Early and Late MCI, Alzheimer) following the ADNI classification criterion. All groups have been balanced with respect to age, sex and years of education. Brain connectivity patterns and its relation with disease progression are accomplished by comparing the control group with the rest of groups, i.e. Control vs EMCI (stage I), Control vs LMCI (stage II) and Control vs AD (stage III).

### 2.6 DTI acquisitions

All subjects in this study had the same ADNI imaging protocol, explained in http://adni.loni.usc.edu/methods/documents/mri-protocols/ and consisting in whole-brain MRI 3T scanners and Diffusion Weighted Images (DWI) images of the axial DTI series. The DTI images were acquired using spin echo pulse sequence echo-planar-imaging (SE-EPI) with the following parameters: TR = 9050.0 ms; TE set to minimum (values ranging from 60 ms till 69 ms); 59 slices with thickness of 2.7 mm with no gap among slices; 128x128 matrix with a FOV of 35.0 cm; with matrix pixels 256x256x2714 and voxel size 1.36x1.36x2.7 mm^3^, flip angle = 90°. A diffusion gradient was applied along 41 non-collinear directions with a b value of 1000 s/mm2. Additionally, one set of images was acquired with no diffusion weighting (b= 0 s/mm2).

### 2.7 Diffusion tensor brain networks

Diffusion tensor brain networks were built following a similar methodology to previous work (Marinazzo *et al.* 2014, Diez *et al.* 2015, Alonso-Montes *et al.* 2015, Amor *et al.* 2015, Diez *et al.* 2017) using FSL (FMRIB Software Library v5.0) and the Diffusion Toolkit. First, all the selected images were downloaded in DICOM and transformed to Nifti format for further analysis. Next, an eddy current correction was applied to overcome the artifacts produced by variation in the gradient field directions, together with the artifacts produced by head movements. Next, using the corrected data, a local fitting of the diffusion tensor was applied to compute the diffusion tensor model for each voxel. Next, a Fiber Assignment by Continuous Tracking (FACT) algorithm was applied (Mori *et al.* 1999). Next, a transformation from the Montreal Neurological Institute (MNI) space to the individual-subject diffusion space was computed and applied to the brain hierarchical atlas (BHA) with M=20 modules, which was shown in (Diez *et al.* 2015) to have the best correspondence between functional and structural modules. This atlas developed by the authors is available to download at http://www.nitrc.org/projects/biocr_hcatlas/. This allowed building 20 x 20 structural connectivity (SC) matrices, each per subject, by counting the number of white matter streamlines connecting all module pairs. Thus, the element matrix *(i,j)* of SC is given by the streamlines number between modules *i* and *j*. As a result, SC is a symmetric matrix, where the connectivity from *i* to *j* is equal to the one from *j* to *i*.

### 2.8 Labelling of anatomical regions

The anatomical representation of the initial 2,514 brain regions existing in BHA was identified by using the Automated Anatomical Labelling (AAL) brain atlas (Tzourio-Mazoyer *et al.* 2002). Therefore, the anatomical identification of the brain regions used in this work followed the labels existing in the AAL atlas.

### 2.9 Cross-group analysis: Multivariate Distance Matrix Regression

The cross-group analysis has been performed using the Multivariate Distance Matrix Regression (MDMR) approach proposed in (Shehzad *et al.* 2014). Connectome-wide association studies are usually performed by means of mass-univariate statistical analyses, in which the association between a phenotypic variable (e.g., the score in a neuropsychological test) with each entry of the brain connectivity matrix is tested across subjects. Such analysis, however, exhibits two main pitfalls: First even at the level of region of interest (ROI) and thus choosing much less regions as voxels, the number of statistical tests entailed is large (Milham 2012), which increases the potential for false positives. On the other hand, studying each brain connectivity matrix entry separately, concurrent contributions from other entries are necessarily ignored (Cole *et al.* 2010). In multivariate methods, instead, the simultaneous contribution of entire sets of brain connectivity entries to a phenotypic variable is evaluated, in a manner that it better captures the concurrent global changes and reduces the number of false positives.

A multivariate distance regression was applied and the variation of distance in connectivity patterns between groups as a response of the Alzheimer’s progression as compared to the Control state was tested. For a fixed brain module *i*, the distance between connectivity patterns of module *i* to the rest of the brain was calculated per pair of subjects (u,v) --by calculating Pearson correlation between connectivity vectors of subject pairs--, thus leading to a distance matrix in the subject space for each module *i* investigated. In particular, the following formula was calculated

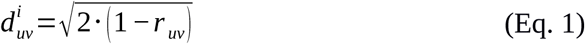

where *r_uv_* is the Pearson correlation between connectivity patterns of *i* for subjects *u* and *v*. After repeating the same procedure for all subjects, as many distance matrices as partition modules (*i*=1,…,20) were obtained. Next, MDMR was applied to perform cross-group analysis as implemented in R (McArtor 2016).

It is important to emphasize that MDMR does not look to how individual modules are locally organized or connected, but to the integration connectivity pattern between those segregated modules to the rest of the brain. Therefore, when group differences were found on a MDMR given module, the connectivity alterations from that module suggests an significant affect to the rest of the brain.

MDMR yielded a pseudo-F estimator (analogous to that F-estimator in standard ANOVA analysis), which addresses significance of disease strength due to between-group variation as compared to within-group variations (McArdle and Anderson 2001). To compare between groups when the regressor variable is categorical (*i.e.* the group label), given a distance matrix, one can calculate the total sum of squares as

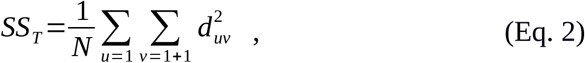

with *N* being the total number of subjects. Notice that, from here on, we will consider 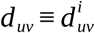. Thus, we got a different *SS_T_* for each module *i*. Similarly, the within-group sum of squares can be written as

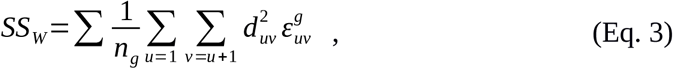

where *n_g_* is the number of subjects per group and 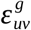 a variable equal to 1 if subjects *u* and *v* belong to group *g* and 0 otherwise. The between-group variation is simply *SS_B_* =*SS_T_ − SS_W_*, which leads to a pseudo-F statistic that reads

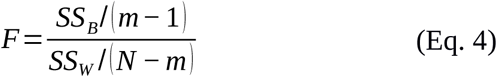

where *m* is the number of groups. As it was acknowledged in (Zapala and Schork 2006), the pseudo-F statistic is not distributed like the usual Fisher's F-distribution under the null hypothesis. Accordingly,we randomly shuffled the subject indices and computed the pseudo-F statistic for each time. A p-value is computed by counting those pseudo F-statistic values from permuted data greater than that from the original data respect to the total number of performed permutations.

Finally, we controlled for type I errors due to the 20 independent statistical performed tests by false discovery rate corrections (Benjamini and Hochberg 1995). Corrected whole-brain connectivity patterns of modules are the ones related to AD progression at the different stages. A schematic overview of the method can be found in Figure 2.

**Figure 2:**
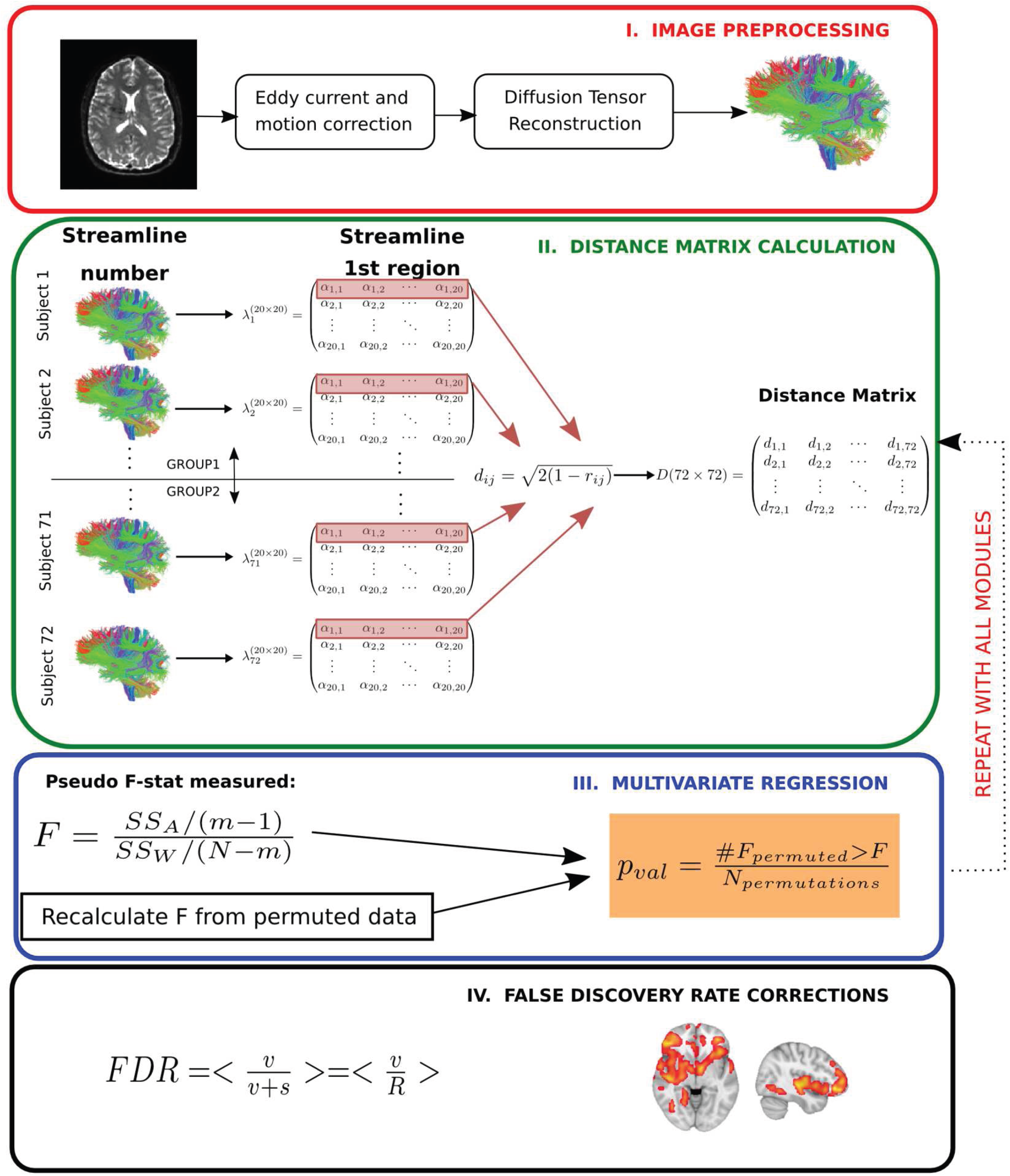
Multivariate distance matrix regression analysis to find differences in brain connectivity patterns across severity progression of AD. In a first step (*Image preprocessing* in a red box), brain images are preprocessed by using standard techniques, mainly eddy current and head motion corrections). Next, diffusion tensor reconstructions are built that allows calculating the tractography for each subject (further details in Methods). In the next step (*Distance matrix calculation* in a green box), first the streamline number connectivity matrix is obtained (here, represented by *λ*), one per subject, corresponding to 20 × 20 entries of values given by *α*. Second, the connectivity patterns of subjects for a given module are used to construct the distance matrix in the subject space by means of Pearson correlation coefficients. Once the distance matrix for a given module is calculated (here, we highlight in red the first row that corresponds to the first module), we test in the third step (*Multivariate regression* in a blue box) whether the variability in distance between different groups is statistically related with disease, for which we compare the observed results with a simulated distribution given by N permutations of the labels. We repeat this operation for every module. We finally apply the fourth step (*False discovery rate corrections* in a black box) to correct for multiple comparisons.

## 3. Results

Results are summarized in Table 3 and modules involved in the disease progression at the group level are shown in Figure 3. See also Table S2 for examples of the different terms participating in the statistical test.

**Table 3:**
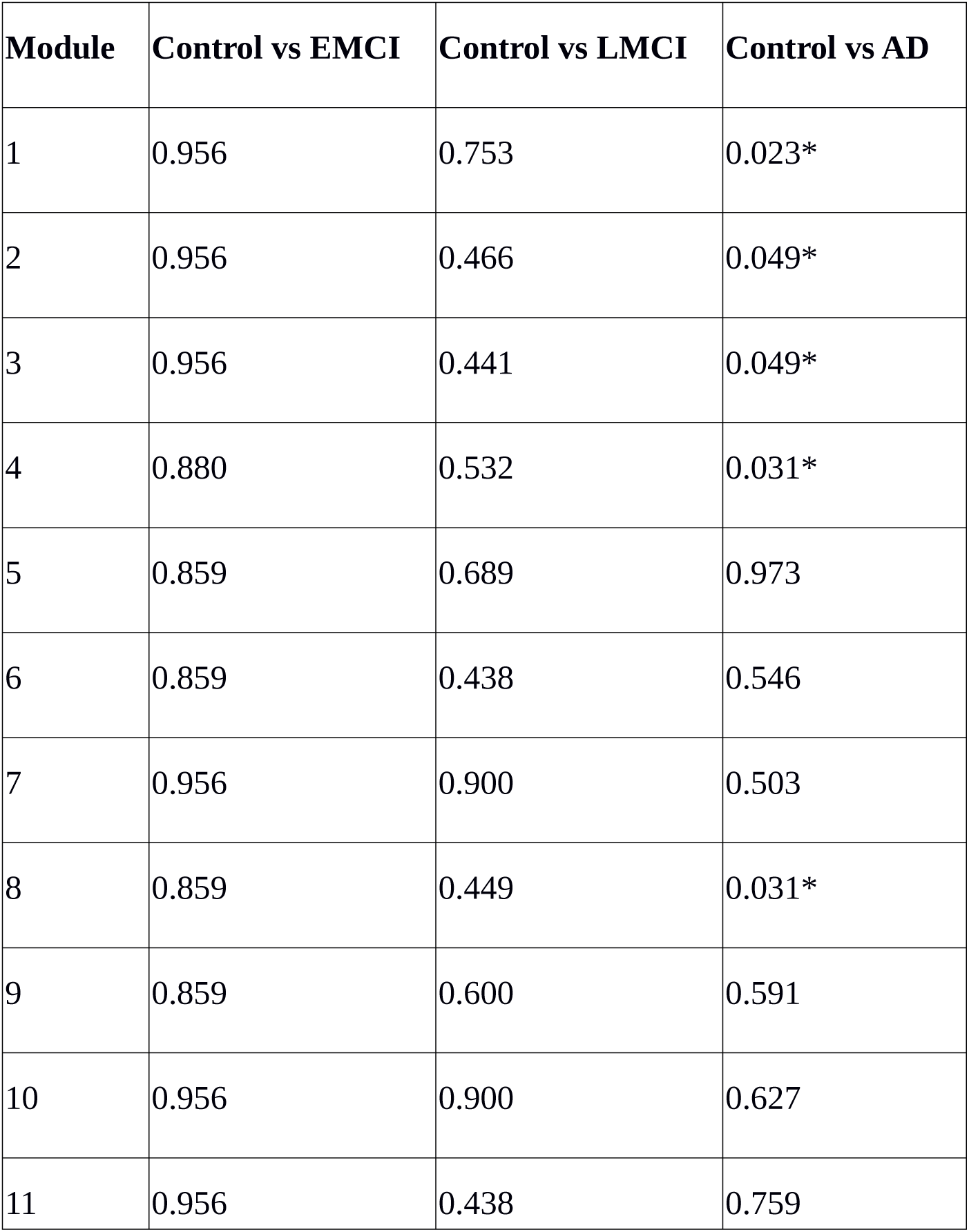
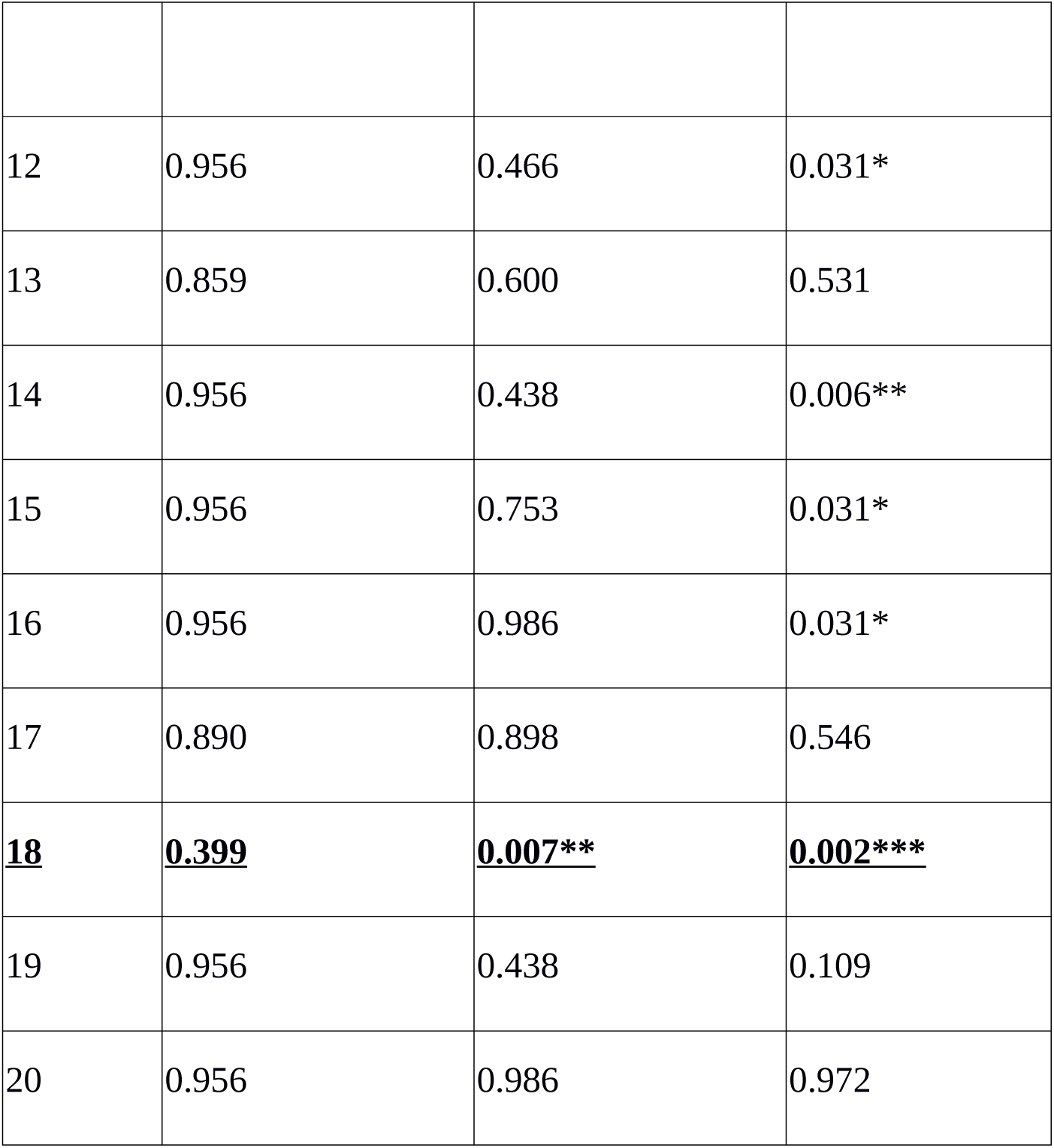
p-values associated to each module from the brain hierarchical atlas. EMCI: Early mild cognitive impairment; LMCI=Late mild cognitive impairment; AD= Alzheimer disease; * 0.01< *p*<0.05; ** 0.005< *p*< 0.01: \*\*\**p*<0.005. Connectivity alterations start in module 18 (marked in black and underlined), and in later stages grow (increasing significance) and extend to a multitude of different other modules.

| Module | Control vs EMCI | Control vs LMCI | Control vs AD |
| --- | --- | --- | --- |
| 1 | 0.956 | 0.753 | 0.023* |
| 2 | 0.956 | 0.466 | 0.049* |
| 3 | 0.956 | 0.441 | 0.049* |
| 4 | 0.880 | 0.532 | 0.031* |
| 5 | 0.859 | 0.689 | 0.973 |
| 6 | 0.859 | 0.438 | 0.546 |
| 7 | 0.956 | 0.900 | 0.503 |
| 8 | 0.859 | 0.449 | 0.031* |
| 9 | 0.859 | 0.600 | 0.591 |
| 10 | 0.956 | 0.900 | 0.627 |
| 11 | 0.956 | 0.438 | 0.759 |
| 12 | 0.956 | 0.466 | 0.031* |
| 13 | 0.859 | 0.600 | 0.531 |
| 14 | 0.956 | 0.438 | 0.006** |
| 15 | 0.956 | 0.753 | 0.031* |
| 16 | 0.956 | 0.986 | 0.031* |
| 17 | 0.890 | 0.898 | 0.546 |
| **<u>18</u>** | **<u>0.399</u>** | **<u>0.007\*\*</u>** | **<u>0.002\*\*\*</u>** |
| 19 | 0.956 | 0.438 | 0.109 |
| 20 | 0.956 | 0.986 | 0.972 |

**Figure 3:**
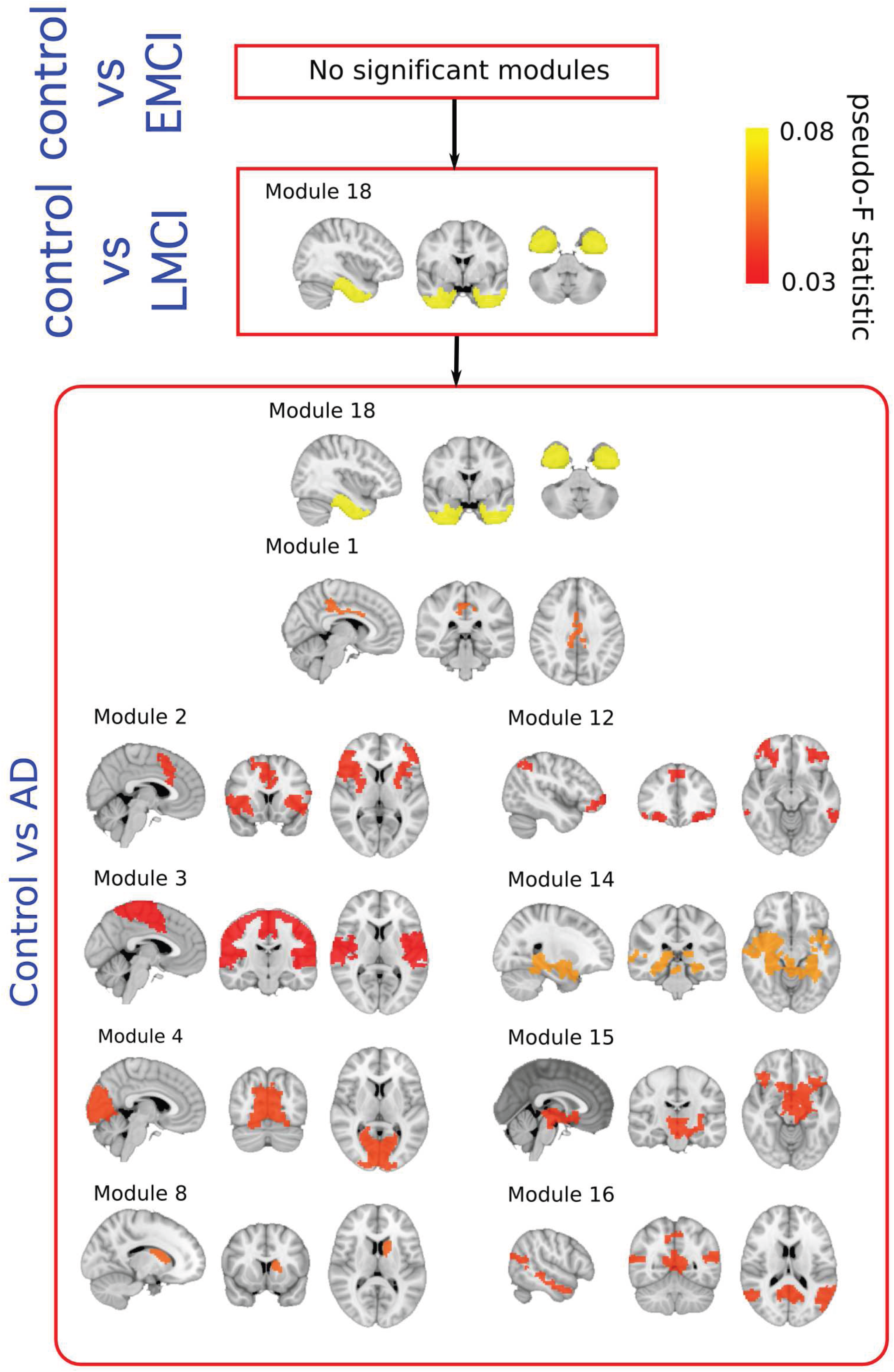
Pseudo F-statistic brain maps across the severity progression of AD. Brain disconnection as disease progresses is quantitatively addressed by looking at the Pseudo F-statistic values of the modules. At first stages (Control vs EMC, top), fibers deterioration is not sufficient to yield significant changes in modules connectivity patterns. In the following stage (Control vs LMCI - middle), the connectivity pattern of module 18, which involves parts of the hippocampus, entorhinal cortex, amygdala and other memory-related areas, disconnects statistically with respect to control (p-val = 0.007). Such connectivity differences are widely spread to the rest of the brain at the final stage (Control vs AD, bottom).

## 3.1 Stage I: Control vs EMCI

A total number of 36 images per each group were selected to perform group comparison. No significant differences were found in terms of module connectivity patterns to the whole brain.

### 3.2 Stage II: Control vs LMCI

A total number of 36 images per each group were selected to perform group comparison. Significant differences were found for the connectivity between the module 18 and the rest of the brain (*p*=0.007). As detailed in (27), the module 18 of the brain hierarchical atlas incorporated part of the hippocampus, amygdala, entorhinal cortex, fusiform gyrus, inferior temporal gyrus, middle temporal gyrus, parahippocampal gyrus and temporal pole.

### 3.3 Stage III: Control vs AD

A total of 36 images per group were selected to perform group comparison. At this stage, significant different connectivity patterns were found in multiple modules existing in BHA:

<u>Module 1</u> (*p*=0.023); including part of the posterior cingulate.

<u>Module 2</u> (*p*=0.049); including part of the putamen, anterior cingulate, rostral pars of the middle frontal gyrus, superior parietal gyrus, supramarginal gyrus, insula, inferior parietal gyrus, precentral gyrus and superior frontal gyrus.

<u>Module 3</u> (*p*=0.049); part of the paracentral lobe, precentral gyrus, postcentral gyrus, precuneus, superior frontal gyrus, superior parietal gyrus, superior temporal gyrus, supramarginal gyrus and insula.

<u>Module 4</u> (*p*=0.031); part of the cuneus, lateral occipital sulcus, lingual gyrus, pericalcarine cortex and precuneus.

<u>Module 8</u> (*p*=0.031); part of the caudate nucleus and putamen.

<u>Module 12</u> (*p*=0.031); part of the inferior parietal gyrus, inferior temporal gyrus, lateral frontal orbital gyrus, pars orbitalis, pars triangularis, rostral pars of the middle frontal gyrus, superior frontal gyrus, caudate nucleus and anterior cingulate.

<u>Module 14</u> (*p*=0.006); part of the thalamus, hippocampus, amygdala, putamen, ventral diencephalon, banks of the superior temporal sulcus, parahippocampal gyrus, superior temporal gyrus, insula, middle temporal gyrus and temporal pole.

<u>Module 15</u> (*p*=0.031); part of the thalamus, putamen, pallidum, brainstem, hippocampus, amygdala, accumbens nucleus, ventral diencephalon, orbital gyrus and insula.

<u>Module 16</u> (*p*=0.031); part of the cerebellum, banks of the superior temporal sulcus, inferior parietal gyrus, cingulate isthmus, middle temporal gyrus, precuneus and superior temporal gyrus.

<u>Module 18</u> (*p*=0.002); see previous 3.2 section for the anatomical description, but notice a reduction in p value from 0.007 (Control vs LMCI) to 0.002 (Control vs AD).

### 3.4 Common affected modules between stages

Connectivity pattern of <u>module 18</u> to the rest of the brain was found at stage II (*p*=0.007) and at stage III (p=0.002), indicating that the further the disease progresses, the greater the connectivity of module 18 is altered to the rest of the brain.

## 4. Discussion

The aim of the present study was to identify differences in brain connectivity patterns between a control group and three pathological groups by disease severity. For this purpose, diffusion tensor brain networks were built allowing determining connectivity differences at three consecutive severity stages: stage I (Control vs EMCI), stage II (Control vs LMCI) and stage III (Control vs AD).

The results showed an absence of significant changes in connectivity patterns in stage I, that is, between patients with early mild cognitive impairment and healthy individuals. The MDMR analysis we have applied finds group differences in the connectivity patterns from different modules to the rest of the brain. Therefore, when observing early mild cognitive impairment, our analysis allows for some possible structural damages to locally occur. This study has shown that even if local alterations exist, they are not capable of producing global inter-module network reorganization/redistribution detectable by the MDMR analysis.

Significant differences were found by the MDMR method in stage II in relation to a network involved with memory (module 18), which includes the hippocampus, amygdala, entorhinal cortex, fusiform gyrus, inferior temporal gyrus, mean temporal gyrus, parahippocampal gyrus and the temporal pole. Strikingly, the change in module 18 connectivity becomes more evident in stage III (i.e., in patients with AD), and *memory* alterations coexist with alterations in a multitude of different modules (modules 1-4, 8, 12, 14-16 and 18), which encompass the default mode network, the sensory-motor network, the medial visual network, frontoparietal regions and subcortical networks (including part of the hippocampus, amygdala and putamen).

The brain connectivity alterations found in this study in stage II might be related to the appearance of several cognitive manifestations, which are typical of AD. For example, many studies have determined the main cognitive impairment in the preclinical phase of AD is episodic memory (Almkvist, 1996, Arnaiz et al., 2003; Albert et al., 2001; Bäckman et al., 2004, 2005; Grober et al., 2008), in which hippocampus; entorhinal cortex and amygdala are involved. Following this line of results, research has found that alterations in the temporal-medial lobe have an affect before AD is even clinically diagnosed (Almkvist, 1996; Bäckman et al., 2004, 2005; Small et al., 1999; Estévez-González et al., 2003; Small et al., 2003). Moreover, research has also shown that the initial neuronal lesions in AD begin in the entorhinal region (included in module 18, therefore, in agreement with our results) with the accumulation of neurofibrillary tangles and neuritic plaques (Gómez-Isla et al., 1996).

Although alterations of the episodic memory are considered the most critical ones at the preclinical phase of AD (Small, et al., 2003; Storandt, 2008) and tasks that measure episodic memory have been shown to be particularly effective at identifying people at risk for developing AD (Elias et al., 2000; Tierney et al., 1996), studies have shown that people with mild cognitive impairment who have altered (in addition to episodic memory) other cognitive areas such as verbal ability (Apostolova et al., 2008; Arnaiz et al et al., 2003; Bäckman et al., 2004, 2005; Joubert et al., 2010), executive functions (Albert et al., 2001; Bäckman et al., 2004, 2005; Dickerson et al., 2007; Grober et al., 2008; Storandt, 2008; Blacker et al., 2007; Rapp et al., 2005), perceptual speed (Bäckman et al., 2005), Visuo-spatial / visuoperceptive skills (Almkvist, 1996, Arnaiz et al., 2003; Bäckman et al., 2004, 2005; Alegret et al., 2009), attention (Bäckman et al., 2005; Rapp et al., 2005), etc. are more likely to convert to AD than those with only memory impairment (Bozoki et al., 2001). As indicated by Bäckman et al. (2004, 2005), a number of different areas in addition to the ones in the temporal-medial lobe are altered prior to the diagnosis of AD (such as the anterior cingulate, temporal sulcus, posterior cingulate, temporoparietal regions, frontal regions and precuneus). This may explain why studies attempting to find cognitive markers of the AD preclinical stage find alterations in other cognitive functions apart from episodic memory.

As the disease progresses, not only the disconnection pattern of module 18 becomes more evident (increasing the distance between AD and controls, Table 3), but such significant changes extend to other brain regions. For example, areas of the hippocampus affected by module 14 are well known to suffer a very severe cognitive degeneration, a fact also confirmed by functional connectivity studies (Zhou *et al.* 2008). The results also indicate a significant connectivity change with temporal medial areas, as revealed by module 16, as shown in Tract Based Spatial Statistics at (Stricker *et al.* 2009, Acosta-Cabronero *et al.* 2010, Salat *et al.* 2010). Similarly to the results of this study, authors of (He *et al.* 2007) demonstrated, through a combined structural and functional analysis, changes in connectivity between the lingual and cuneus, by using only structural connectivity data.

The results of the present study indicate a significant change in the connectivity from the entire brain to the areas provided by module 4, mainly associated to visual function. A decrease in virtual capacity in AD is well known, especially in those areas involving movement blindness, depth perception, color perception and contrast sensitivity (Whittaker *et al.* 2002). Again, this damage expansion to other brain regions also agrees with the extent and worsening of cognitive aspects (e.g., memory, attention, language; Weintraub *et al.* 2012) and neurobehavioral problems (e.g. personality changes, anxiety, depression, agitation, hallucinations; Chung & Cummings, 2000, Bassiony et al., 2000, Senanarong et al., 2003) of patients with AD.

Previous studies have analyzed the connectivity differences from tensor diffusion networks in AD and have found significant alterations in the inferior longitudinal fasciculus for patients at risk of AD (Smith *et al.* 2010), which could correspond to LMCI. Similarly, a voxel-based analysis in (Honea *et al.* 2009) showed a significant decrease in FA for fibers connecting the parahippocampal gyrus. In addition, patients diagnosed in the early stages of AD (corresponding to early or late mild cognitive impairment in this study) had a significant reduction in white matter in the upper longitudinal fasciculus, which also connects part of module 18 in the brain hierarchical atlas with the frontal lobe (Rose *et al.* 2000). The authors (Hanyu *et al.* 1998) found significant changes in apparent diffusion coefficients and diffusion anisotropy in patients with recent progressive cognitive impairment, suggesting an early decrease in temporal fiber density, a region included in the module 18, therefore in concordance to our results.

### A different comparison between pathological groups

By defining disease progression across three stages, I (control vs EMCI), II (control vs LMCI) and III (control vs AD), we have found progressive variations in connectivity patterns that start in a module clearly associated to memory function (including part of the hippocampus, amygdala, entorhinal cortex, fusiform gyrus, inferior and middle temporal gyrus, parahippocampal gyrus and temporal pole) and later on, alterations are found widespread along the entire brain. Therefore, it is important to emphasize that we have defined disease progression by comparing each pathological group with respect to the control group. A different possibility for assessing connectivity variations is to perform comparisons between pathological groups, i.e., EMCI vs LMCI, LMCI vs AD, EMCI vs AD. For the two comparisons EMCI vs LMCI and LMCI vs AD, none of the module showed differences in connectivity patterns (Table S3). However, the EMCI vs AD comparison showed differences in modules 2,3,4,14 and 16.

The reason why our strategy of defining disease progression with respect to the control group found differences in module 18 at the beginning of the progression is due to the fact that the within-group distance contribution of the control group is smaller than the corresponding one in any of the pathological groups. In particular, we calculated the sum of distances squared (defined in Eq. 1) between pairs of subjects of connectivity between module 18 and the rest of the brain and obtained values of 62 (control), 76 (EMCI), 83 (LMCI) and 82 (AD). In other words, the tensor-diffusion connectivity values of module 18 are more homogeneous between subjects within the control group as compared to subjects within any other pathological group, what makes our strategy to successfully detect differences in the connectivity pattern of module 18 at the early stages of disease progression.

### Implications

In recent years a great deal of emphasis has been placed on early AD detection (Albert et al., 2001); from looking for pharmacological or non-pharmacological treatments to help delay the age of onset disease and to slow down the clinical disease progression. Similar to other studies, these results provide (by looking to diffusion tensor brain networks) that the earliest detection in connectivity patterns affecting globally the rest of the brain starts in a network mainly encompassing memory function.

On the other hand, identifying brain connectivity patterns in patients who have not yet developed AD might shed some light in determining how these connectivity patterns evolve as time goes on. In addition, it will be possible to associate connectivity patterns with clinical patient's variations existing at each disease stage. This might help better understand the relationship between deterioration in brain functioning and clinical patient's characteristics.

### Limitations

The results of the present study should be interpreted in light of the following limitations. First, it is a cross-sectional study with different groups of patients in each experimental group and with a small sample size, so future studies should try to extend to bigger cohorts and follow the same group of people over time as the disease progresses. Second, the patients included in the study have a probable AD, which means that the definitive diagnosis of AD can only be performed post-mortem (Fearing et al., 2007). The use of patients with familiar AD could help to know in depth the evolution of the disease and the changes in cerebral connectivity from many years back to its onset. Third, there are a number of risk factors associated with the decline of mild cognitive impairment which can affect brain connectivity such as advanced diabetes, symptomatology depressive disorder, hypertension, hypotension, obesity, history of traumatic brain injury and APOE genotype, that have not been taken into account in this study. Future studies should take into account the possible influence of these variables on the processes of cerebral connectivity.

### Summary

In conclusion, the results obtained from this study applying a multivariate method to diffusion tensor connectivity networks across AD severity progression, are in line with the evolution of AD from both the neuropathological and neuropsychological points of view. That is, first alterations occur in the connectivity of regions of the middle temporal lobe (hippocampus and entorhinal), which coincides with the first symptoms of altered episodic memory in the preclinical stage and in mild cognitive impairment. As the disease progresses, the brain damage and its disconnection of these regions become more evident and expands to other areas, which coincides with the expansion and/or worsening of other cognitive functions and neurobehavioral aspects seen in the individuals with AD. Future developments will deal with the application of the same methodology to longitudinal data, a mandatory step to confirm our results.

## Author Contributions

JR, CAM and ID had equal first-author contribution; JR, CAM and ID analyzed the data and made the figures; LOL and JCAL connected results to cognitive deficits in AD; LR, IE, BM, PB, MF, JCAL, SS and JMC designed the research; all the authors wrote the manuscript and agreed in its submission; SS and JMC had equal last author contributions

## Acknowledgements

SS acknowledges financial support from Bizkaia Talent and European Commission through COFUND with the research project BRAhMS – Brain Aura Mathematical Simulation– (AYD-000-285). JR acknowledges financial support from the Minister of Education, Language Policy and Culture (Basque Government) under Doctoral Research Staff Improvement Programme. JMC and JCAL acknowledge financial support from Ikerbasque (The Basque Foundation for Science) and from Ministerio Economia, Industria y Competitividad (Spain) and FEDER (grant DPI2016-79874-R). PB acknowledges financial support from Ikerbasque and from the Ministerio Economia, Industria y Competitividad (Spain) and FEDER (grant SAF2015-69484-R).

Data collection and sharing for this project was funded by the Alzheimer's Disease Neuroimaging Initiative (ADNI) National Institutes of Health grant U01 AG024904. ADNI is funded by the National Institute on Aging, the National Institute of Biomedical Imaging and Bioengineering, and through generous contributions from the following: Abbott, AstraZeneca, AB, Amorfix, Bayer Schering Pharma AG, Bio-clinica Inc., Biogen Idec, Bristol-Myers Squibb, Eisai Global Clinical Development, Elan Corporation, Genentech, GE Healthcare, Innogenetics, IXICO, Janssen Alzheimer Immunotherapy, Johnson and Johnson, Eli Lilly and Co., Medpace, Inc., Merck and Co., Inc., Meso Scale Diagnostic, & LLC, Novartis AG, Pfizer Inc, F. Hoffman-La Roche, Servier, Synarc, Inc., and Takeda Pharmaceuticals, as well as non-profit partners the Alzheimer’s Association and Alzheimer’s Drug Discovery Foundation, with participation from the U.S. Food and Drug Administration. Private sector contributions to ADNI are facilitated by the Foundation for the National Institutes of Health (www.fnih.org). The grantee organization is the Northern California Institute for Research and Education, and the study is coordinated by the Alzheimer's Disease Cooperative Study at the University of California, San Diego. DATA are disseminated by the Laboratory for Neuro Imaging at the University of California, Los Angeles. This research was also supported by NIH grants P30AG01012 9, K01 AG0305 14, and the Dana Foundation.

## brain connectivity

**Table S1:**
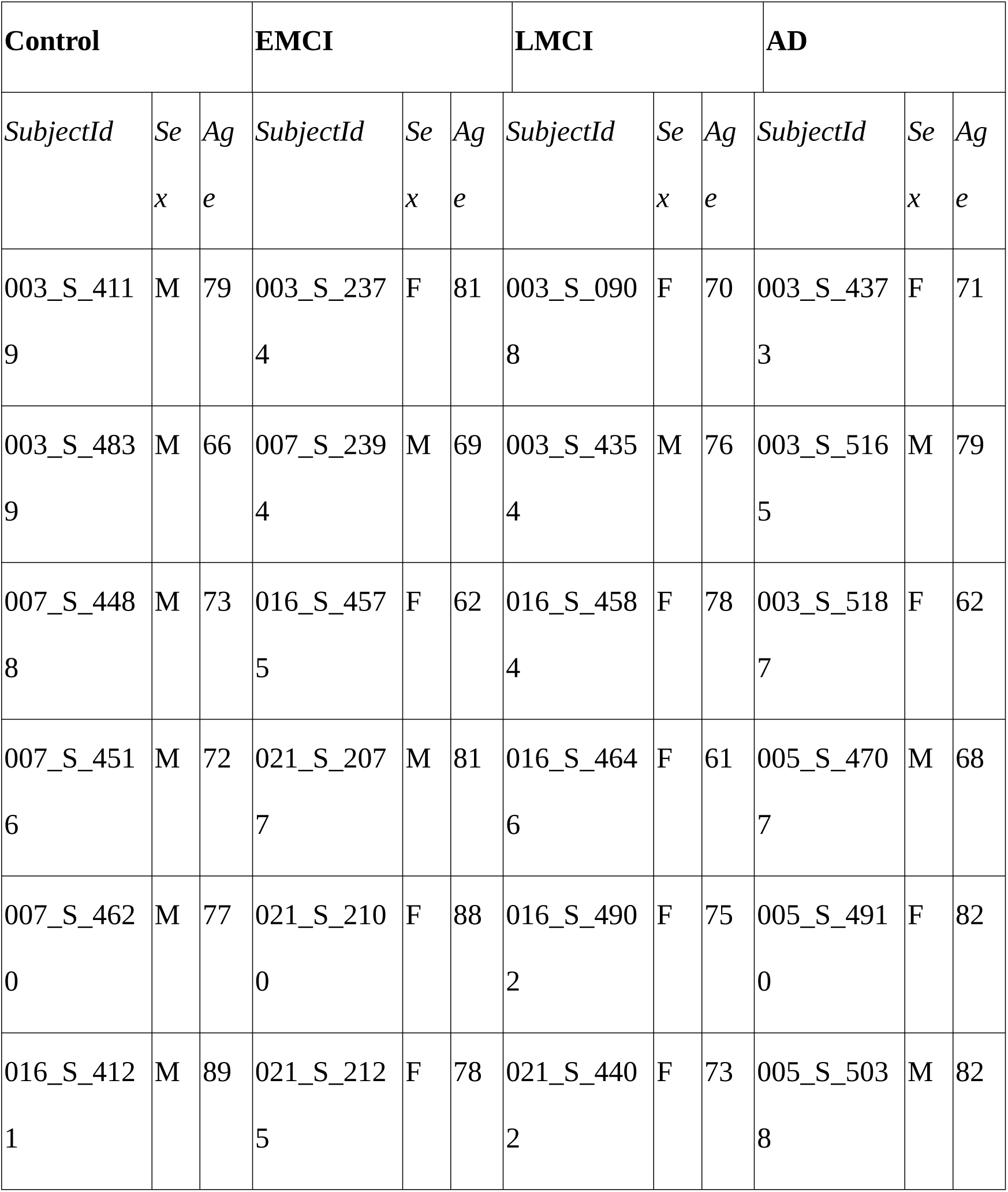
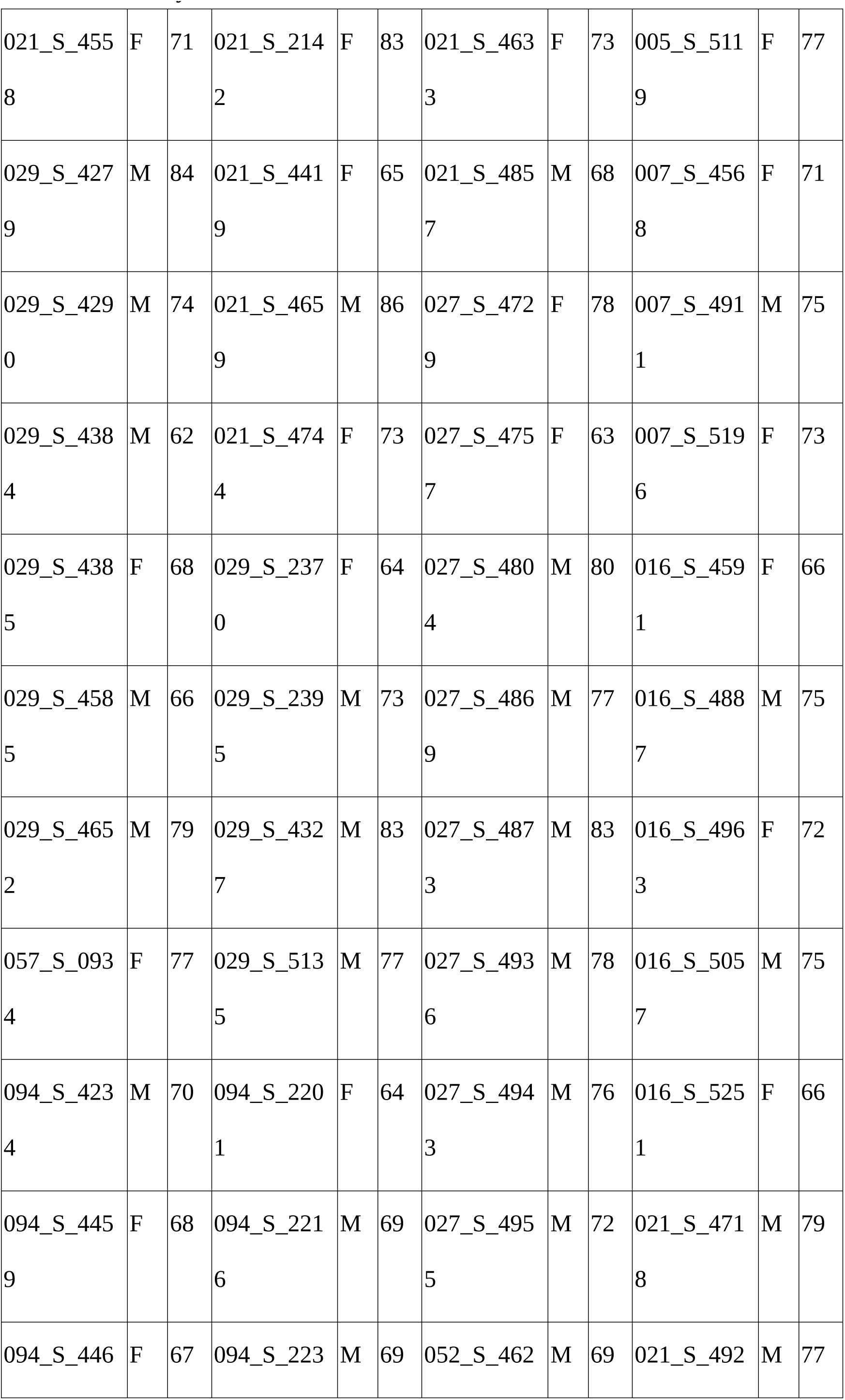
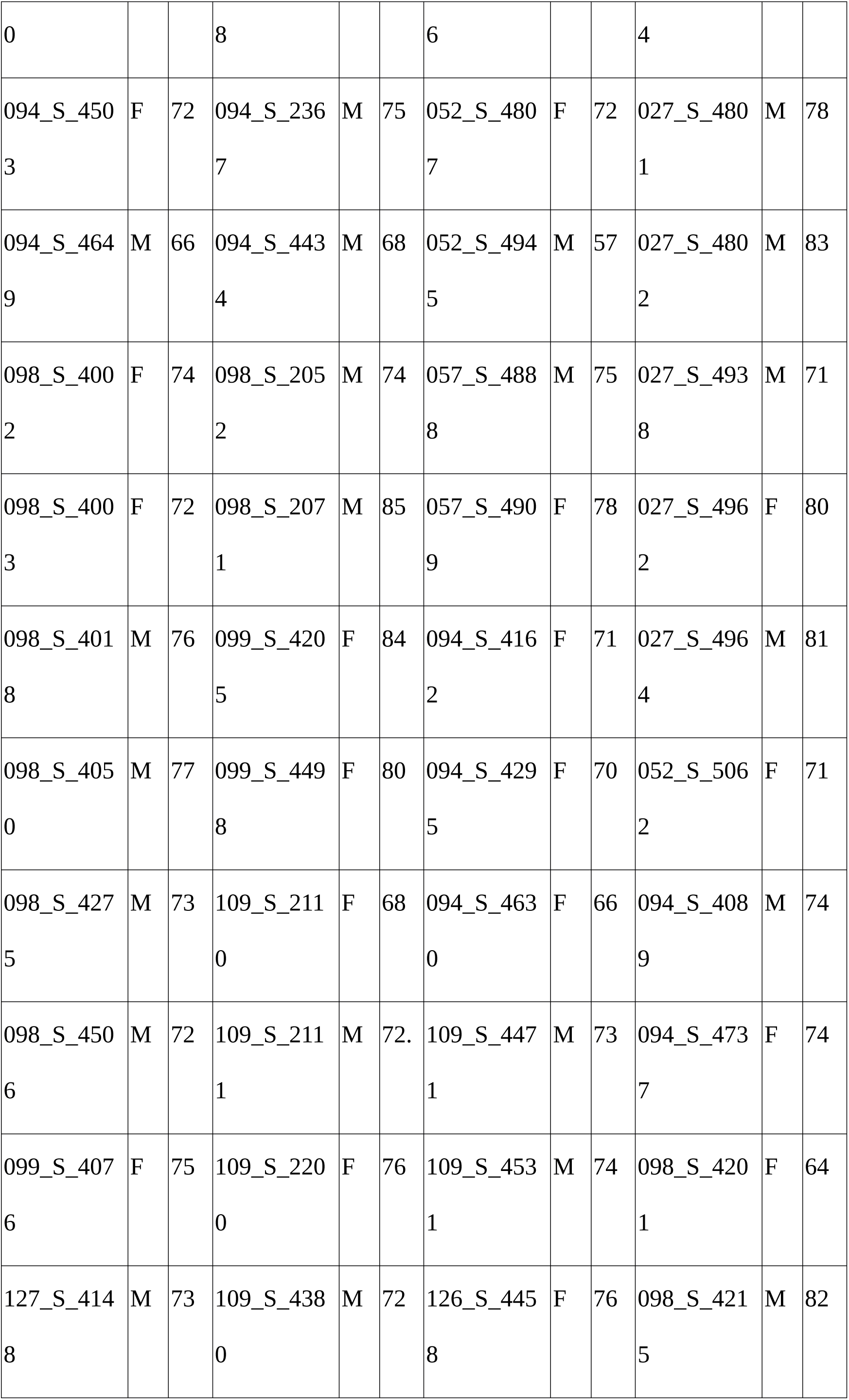
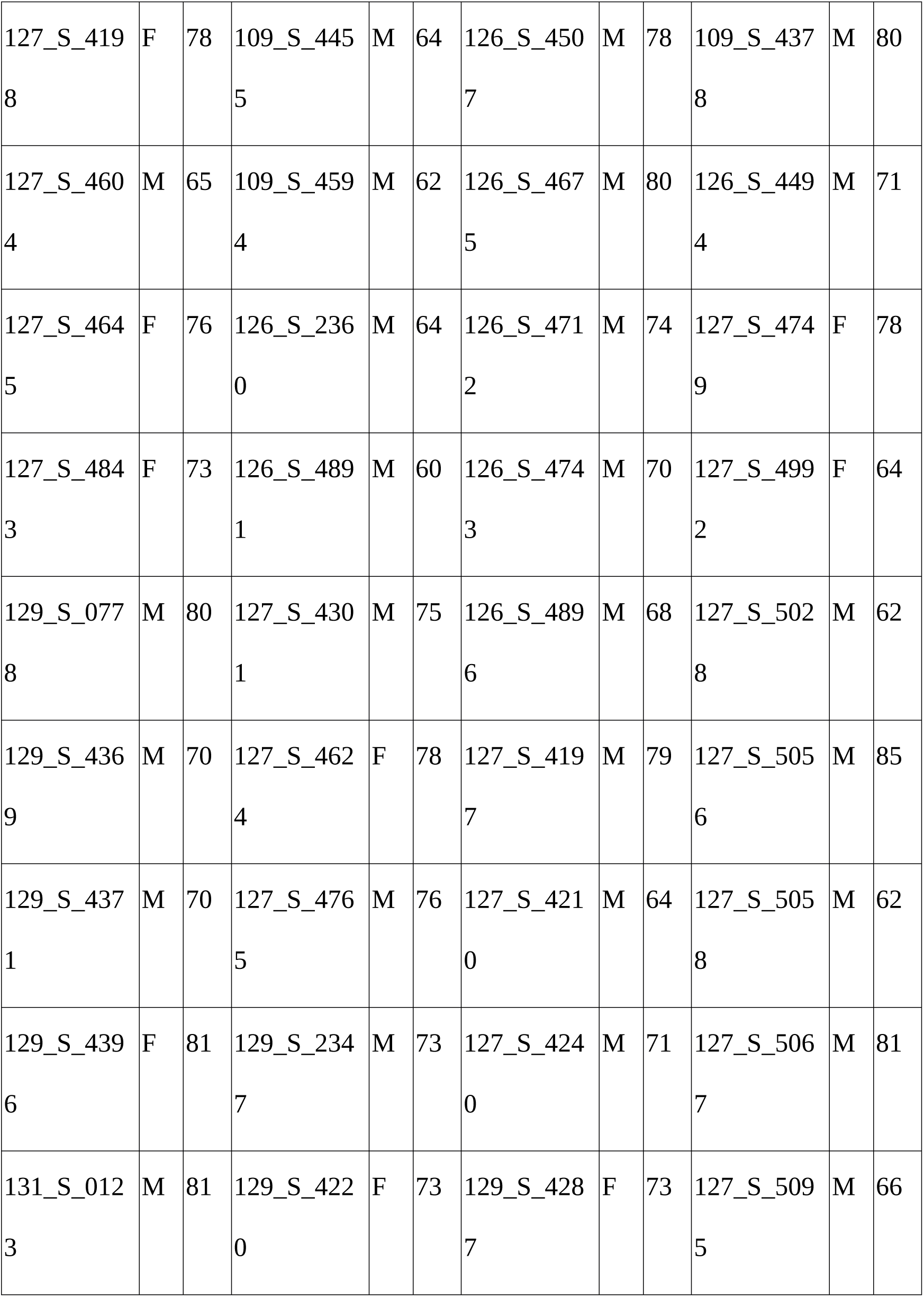
ADNI subjects within each group EMCI: Early mild cognitive impairment; LMCI=Late mild cognitive impairment; AD= Alzheimer disease; M=Male, F=Female.

**Table S2:** Examples of pseudo F-statistics, between-group and within-group sum of squares. Three diferent situations: Node 10, that does not provide any signiicant change in pattern connectivity; Node 16, signiicantly diferent in stage III; and Node 18, with pattern connectivity signiicantly diferent in stages II and III. EMCI: Early mild cognitive impairment; LMCI=Late mild cognitive impairment; AD= Alzheimer disease

| Module | Control vs EMCI |  |  | Control vs LMCI |  |  | Control vs AD |  |  |
| --- | --- | --- | --- | --- | --- | --- | --- | --- | --- |
|  | *F* | SSA | SSW | *F* | SSA | SSW | *F* | SSA | SSW |
| 10 | 0.527 | 0.008 | 1.028 | 0.630 | 0.008 | 0.940 | 0.894 | 0.012 | 0.941 |
| 16 | 0.380 | 0.019 | 3.457 | 0.285 | 0.014 | 3.404 | 3.410 | 0.173 | 3.550 |
| 18 | 3.057 | 0.024 | 0.558 | 5.854 | 0.049 | 0.595 | 6.018 | 0.051 | 0.588 |

**Table S3:**
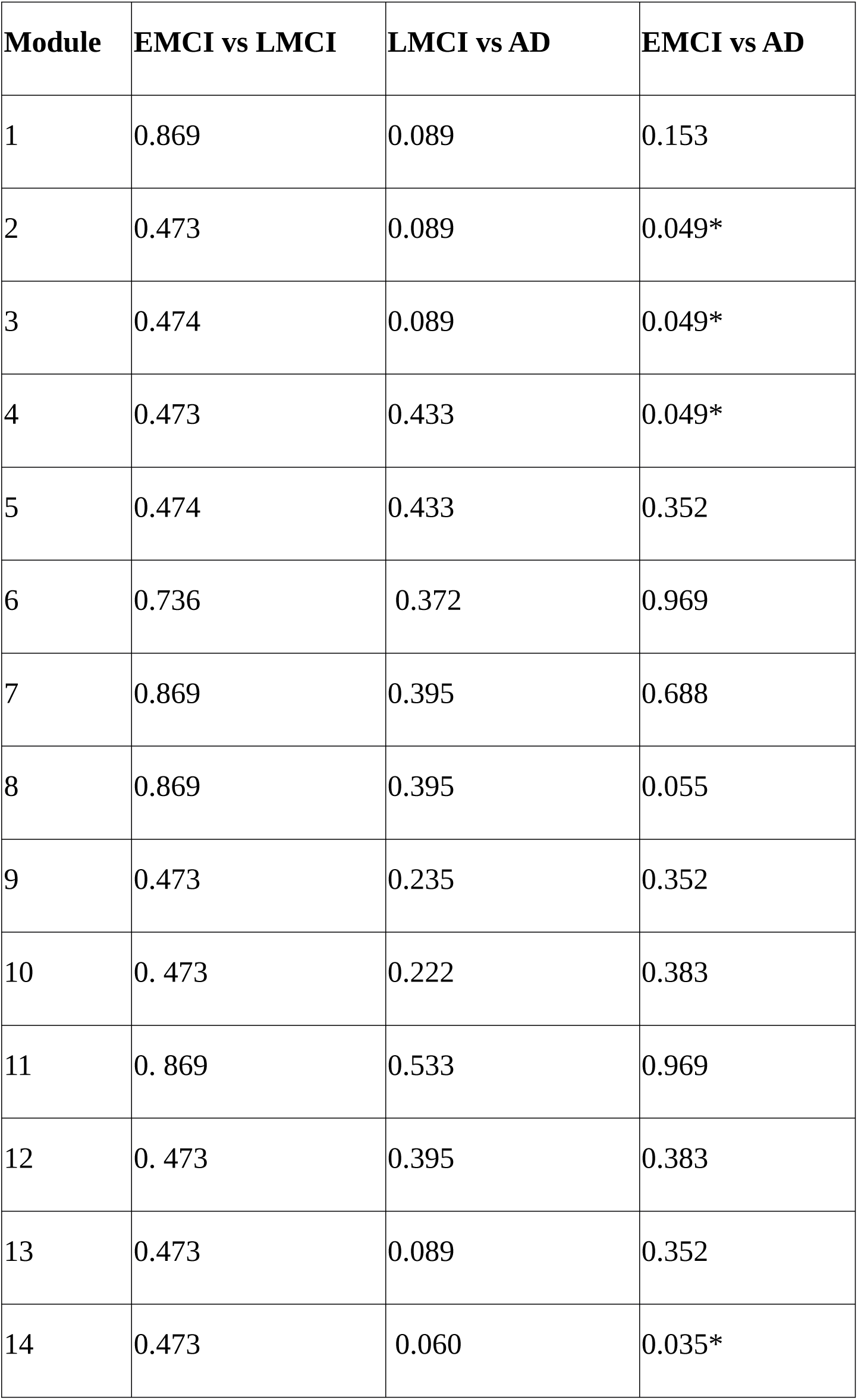
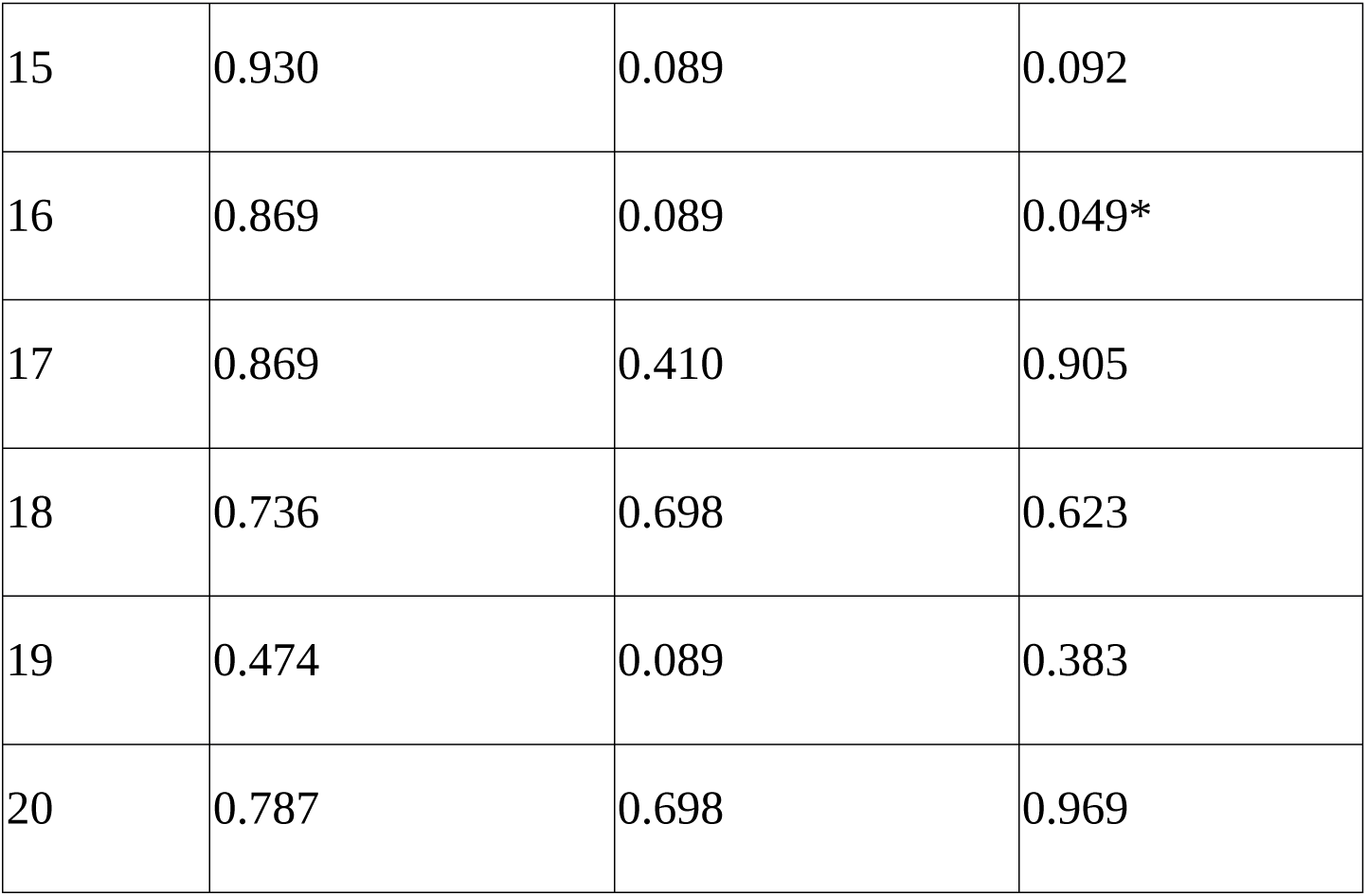
Group comparison not involving the healthy control group. EMCI: Early mild cognitive impairment; LMCI=Late mild cognitive impairment; AD= Alzheimer disease * 0.01<*p*<0.05; ** 0.005<*p*<0.01: *** *p*<0.005.

| Module | EMCI vs LMCI | LMCI vs AD | EMCI vs AD |
| --- | --- | --- | --- |
| 1 | 0.869 | 0.089 | 0.153 |
| 2 | 0.473 | 0.089 | 0.049* |
| 3 | 0.474 | 0.089 | 0.049* |
| 4 | 0.473 | 0.433 | 0.049* |
| 5 | 0.474 | 0.433 | 0.352 |
| 6 | 0.736 | 0.372 | 0.969 |
| 7 | 0.869 | 0.395 | 0.688 |
| 8 | 0.869 | 0.395 | 0.055 |
| 9 | 0.473 | 0.235 | 0.352 |
| 10 | 0.473 | 0.222 | 0.383 |
| 11 | 0.869 | 0.533 | 0.969 |
| 12 | 0.473 | 0.395 | 0.383 |
| 13 | 0.473 | 0.089 | 0.352 |
| 14 | 0.473 | 0.060 | 0.035* |
| 15 | 0.930 | 0.089 | 0.092 |
| 16 | 0.869 | 0.089 | 0.049* |
| 17 | 0.869 | 0.410 | 0.905 |
| 18 | 0.736 | 0.698 | 0.623 |
| 19 | 0.474 | 0.089 | 0.383 |
| 20 | 0.787 | 0.698 | 0.969 |

